# Trait lability as a predictor of diversification dynamics in flowering plants

**DOI:** 10.1101/2024.06.03.597046

**Authors:** James D Boyko, Thais Vasconcelos

## Abstract

Rates of diversification differ between angiosperm lineages. To date, attempts to explain this heterogeneity have focused on the potential correlation between speciation and extinction rates and particular key traits. However, an often-overlooked explanation is that evolutionary lability, here defined as the rates of trait change, may be a better predictor of speciation and extinction rate heterogeneity than the observed traits themselves. Here, we show how this can be tested by using hidden Markov models (HMMs), which allow for several rate classes associated with speciation, extinction, and transition between trait states across a phylogeny. Using a phylogenetic dataset of 13 angiosperm clades including 10,474 species, we show that higher rates of change between open and closed-canopy biomes is consistently associated with higher lineage turnover rates (speciation + extinction rates) across clades. We demonstrate how HMMs can be leveraged in ways that go beyond their conventional use as null models in diversification analyses, and that comparing different rate classes can unveil novel patterns of biological interest. These patterns result in a shift in focus from static traits to dynamic evolutionary processes and may provide a more comprehensive understanding into how biodiversity is generated and maintained, in angiosperms and other organisms.

## Introduction

It is widely accepted that rates of speciation and extinction differ between clades. Nonetheless, the cause of this heterogeneity remains poorly understood despite the consequences being immediately apparent. Angiosperms, or flowering plants, are by far the most diverse clade of green plants alive today with c. 300,000 described species [1]. But less appreciated is that the same degree of disparity in species richness between angiosperms and other green plant clades is repeatedly observed among clades within angiosperms. For example, there are ten times more species of daisies (Asteraceae) than those of primroses (Primulaceae s.s.) and nine times more species of grasses (Poaceae) than sedges (Cyperaceae) [2]. This disparity can be observed at all taxonomic levels and between clades which have had the same amount of time for species accumulation [3].

Differences in clade richness are mainly explained by differences in rates of speciation and extinction. To date, most explanations of clade specific diversification patterns have relied on discovering innovative traits, novel landscapes, or both (“key opportunities”) which correlate with diversification rates [4–7]. Such events are theorized to correlate with increased or decreased chances of speciation or extinction due to the opportunity for population isolation, differentiation, or resilience against extinction [4–7]. Though this search has been fruitful, there has been a lack of consistency in which traits are affecting diversification dynamics in general. In fact, despite hundreds of studies describing a relationship between traits, biogeography, and diversification (reviewed in [8]), studies which have utilized modern, more appropriate, null models (e.g. [9]) have often found little support for character dependent diversification. Even in cases where there is evidence for character dependent diversification, the characters are not consistent drivers of diversification across multiple clades [10].

This lack of generalities suggests that alternative explanations for observed heterogeneity in rates of diversification should be sought. One potential venue that has been suggested is that rates of speciation and extinction correlate with rates of trait change – that is, trait lability [6,8,11]. The botanist G. Ledyard Stebbins was perhaps one of the first to explicitly suggest this possibility in flowering plants [12]. He proposed that the high capacity of angiosperms to adapt to environmental change was the key to their rich diversity. His mechanism was that long term climatic instability led to cyclical changes in the location of climatic zones [13]. For lineages in these unstable regions, the constant fluctuation in the climatic setting would be responsible for continuous allopatry and adaptive pressure for phenotypic change, as well as leading to many ephemeral species and subsequent extinction [12,14]. Furthermore, it is expected that, although observed patterns at the macroevolutionary scale suggest that plant lineages tend to track their preferred environments over time [15,16], when change is rapid, tracking is likely to be imperfect, and phenotypic change or extinction will follow [12,14]. In other words, the continuous movement of lineages between biome types would increase probabilities of both speciation and extinction over time, i.e. evolutionary turnover (sensu [17]).

Another possible reason for the lack of consensus on the drivers of diversification is methodological. For instance, early work quantified diversification rates across angiosperm clades revealing the uneven distribution of species richness without testing potential explanatory factors [3]. As the field advanced, more complex models emerged, with State-dependent Speciation and Extinction (SSE) models becoming particularly popular for testing hypotheses about the link between specific traits and diversification rates [18]. However, this initial enthusiasm was tempered by the growing recognition that standard SSE models were prone to high rates of false positives [19]. This led to the development of new “null” SSE models which incorporate “hidden” variation by applying a hidden Markov model (HMM) framework states and account for rate heterogeneity that is not associated with the observed trait of interest [9]. Despite these methodological improvements, studies have often found a lack of support for trait-dependent diversification when these more robust models are applied [10]. This suggests that while hidden Markov models are useful for avoiding spurious correlations, they also highlight the complexity of diversification dynamics.

Here, we test the hypothesis that rates of transition between open-canopy biomes and closed canopy biomes positively correlate with turnover and net-diversification rates across many angiosperm clades. To this end, we use a modeling framework that combines properties of HMMs [20] and state-speciation and extinction (SSE) models [9,18] to allow for both jointly estimating transition and diversification dynamics in a clade and accounting for heterogeneity in those rates at different parts of the phylogenetic tree, i.e. different rate classes. For our dataset, we sample 13 flowering plant clades, which combined span about 10,474 species where 5,257 species are found in closed-canopy, 3,305 in open-canopy and 1,912 are widespread across both biome types. For each clade, we build hidden Markov SSE models and interpret parameter estimates using non-parametric sign tests. In this way, we are able to explicitly test whether rate classes with faster transition rates between biomes also tend to be those with faster rates of speciation and extinction.

## Methods

### Dataset assembly

The criterion to select the 13 clades used in this study was to pick clades that had a minimum of 250 tips, with at least 0.4 of sampling fraction for the ingroup, i.e. inclusion of at least 40% of the species diversity assigned to the corresponding clades. Although this is a somewhat arbitrary threshold, it has been demonstrated that very low sampling fraction often lead to inaccurate parameter estimates in trait dependent diversification analyses [21,22]; and trees with less than 300 tips are unlikely to have the necessary information to properly fit complex SSE models [23] and to present rate class heterogeneity (see details about rate classes below). We kept our thresholds slightly more inclusive than these (i.e., including clades with as low as 0.4 sampling fraction in Solanaceae and 267 tips in Onagraceae) to retain a minimum diversity of clades across the angiosperm tree of life. We also focused on trees produced by taxonomists of each group, since they are more careful about voucher identification and choice of molecular markers used in reconstructions (e.g. [24]). We avoided using large trees inferred by genbank scraping (e.g. [25]) because of issues with uneven sampling across the tree. In that way, we combined strengths of the large sample approach in which to observe generalities from with the care for accuracy that comes from small trees (see debate between [26] and [27]). Finally, we also excluded trees that were not reciprocally monophyletic, keeping only the ones with larger sampling and/or more tips when the same clade appears in more than one tree (e.g. we excluded the *Acacia* phylogeny of [28] and kept the phylogeny of Mimosoid legumes of [29], which includes *Acacia*). Tree root ages ranged from 9.84 (NeoAstragalus; [30]) to 105.71 (Fagales; [31]) million years to the root and from 267 (Onagraceae; [32]) to 2,202 (Arecaceae, [33]) tips. To score the main biome type for each species, we first standardized all tip names according to the GBIF taxonomic backbone using the R package taxize [34] and removed outgroups following the original publications of each phylogeny. We also pruned trees to remove tips of species with multiple entries in some phylogenies, leaving only one tip per species.

We then downloaded all the 41,496,903 occurrence points for the originally 11,593 species sampled in the 13 trees from GBIF [35–47], using POWO [48] taxonomy and geographical information to filter non-native distributions and distribution inaccuracies. Because GBIF is prone to sampling bias, we also thinned occurrence points to five representative occurrence points per 1×1 grid cell for all species before further analyses (see [49] for detailed methodology). Our filtering decreased the number of species with trustworthy information to 10,474. The remaining 2,264,540 points after filtering were overlaid on the World Wildlife Fund (WWF) map of terrestrial biomes [50]. Biomes are generally defined as a combination of climatic factors that drive convergent and/or parallel evolution in certain plant traits, leading to characteristic vegetation types (or physiognomies) under specific climatic conditions [51].

Following Stebbins’ hypothesis, we binarized the original 13 terrestrial biomes into closed or open-canopy, dividing them along a precipitation gradient. Occurrence points placed on biomes where precipitation is higher (“Tropical & Subtropical Moist Broadleaf Forests”, “Tropical & Subtropical Dry Broadleaf Forests”, “Tropical & Subtropical Coniferous Forests”, “Temperate Broadleaf & Mixed Forests”, “Temperate Conifer Forests”, and “Boreal Forests/Taiga”) were scored as closed-canopy biomes. Those occurring in areas of lower precipitation (“Tropical & Subtropical Grasslands, Savannas & Shrubland”, “Temperate Grasslands, Savannas & Shrublands”, “Flooded Grasslands & Savannas”, “Montane Grasslands & Shrublands”, “Tundra”,”Deserts & Xeric Shrublands”, “Mediterranean Forests, and Woodlands & Scrub”) as well as “Mangroves” were scored as open-canopy biomes. Of course, that does not mean that closed-canopy biomes only have closed-canopy habitats, as open-canopy vegetations are often found within closed-canopy biomes and vice versa (e.g. prairies in temperate rainforests, riverine forests in tropical savannas). It means closed canopy biomes are dominated by closed-canopy vegetations and open-canopy biomes by open-canopy vegetations. Species were scored as occurring in closed or open-canopy biomes when at least 25% of their occurrence points were found in that biome type. If at least 25% of their occurrence points were found in both biome types, then the species was scored as widespread. All datasets and R scripts used in the curation process are available at [10.5281/zenodo.17401294].

### Diversification and hidden state modeling

For each of the 13 angiosperm clades we use the R-package ‘hisse’ [52] to test for correlations between diversification dynamics and rate of biome shift. We fit a set of 36 models: 4 without hidden states (one rate class) and 32 with hidden states (two rate classes). The basic discrete character model of evolution was an all-rates-different model, in which transitions between open and closed-canopy biome could only occur through an intermediate widespread state (Figure 1A). Models with two rate classes were categorized into two types of discrete character models. The first type allows for discrete character rate heterogeneity by permitting transition rates to differ between rate classes. The second type has transition rates that are fixed to be equal between rate classes. For the MuHiSSE model, in addition to trait evolution, we must define the structure of the diversification dynamics in terms of extinction fraction (extinction/speciation) and turnover (speciation+extinction).

**Figure 1.**
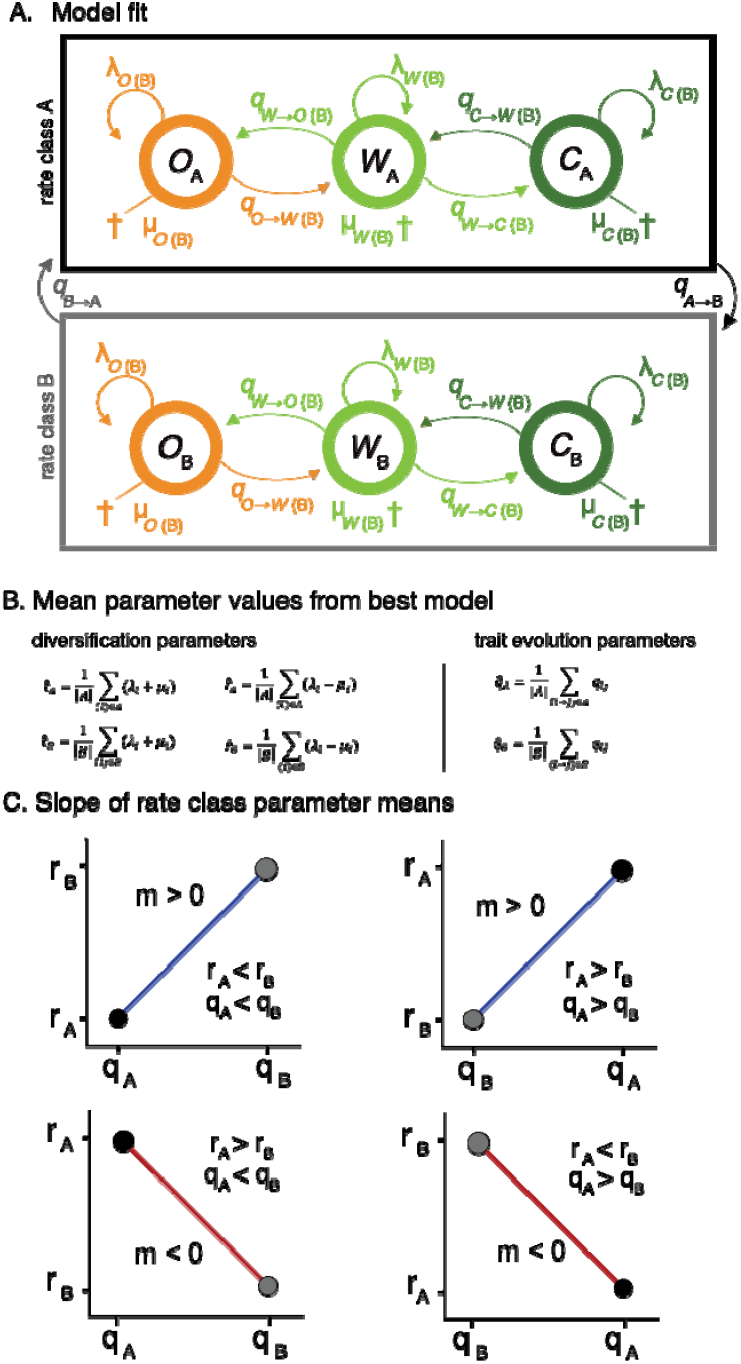
Modeling framework for testing the association between biome shifting and diversification dynamics. (A) A visual representation of the MuHiSSE model, which incorporates three observed biome states (O = Open-canopy, W = Widespread, C = Closed-canopy) and two unobserved “hidden” rate classes (A and B). Within each state and class, the model estimates rates of speciation (λ), extinction (µ), and biome transition (q). (B) Illustration of the parameter extraction from the best-fit model for a single clade. For each of the two hidden rate classes, we extracted the mean biome transition rate and the mean evolutionary turnover rate (speciation + extinction). (C) Each clade is treated as an independent data point testing the association between biome transition rate and turnover. A “positive relationship” (blue lines) is recorded if the hidden rate class with higher transition rates also has higher turnover. A “negative relationship” (red lines) is recorded if it has lower turnover.

For single rate class models, parameters are fixed to be equal for all observed states (no trait-dependent diversification) or all are set to be different (biome dependent diversification). For two rate class models, we explore 4 different parameterizations: (1) no observed trait-dependent diversification and no hidden trait-dependent diversification, (2) observed trait-dependent diversification, but no hidden trait-dependent diversification, (3) no observed trait-dependent diversification, but hidden trait-dependent diversification, and (4) both observed trait-dependent diversification and hidden trait-dependent diversification. We examined all possible combinations of the parameterizations resulting in 36 model structures, and the best fitting model for each clade was used to estimate the transition, speciation and extinction parameters for that clade (Tables S1). Estimates of sampling fraction, a fixed parameter in the model, were gathered from the literature of each clade (Table S2).

### Parameter interpretation and analyses

Following model selection, we analyzed the parameter estimates from the best-fit model for each clade to test for a consistent association between the rate of biome shifts and evolutionary turnover. For each clade whose best-fit model included two distinct rate classes (e.g., rate class ‘A’ and rate class ‘B’), we extracted the mean biome transition rate, the mean turnover rate (speciation + extinction), and mean net diversification rate (speciation - extinction) separately for both rate classes (Figure 1B). A positive relationship was recorded if the rate class with higher biome transition rates also had higher turnover (or net diversification), while a negative relationship was recorded if it had lower rates (Figure 1C). To evaluate if a positive association occurred more frequently than expected by chance across all clades, we conducted separate non-parametric sign tests for turnover and net diversification.

A non-parametric approach is particularly appropriate for this analysis. It treats each clade as an independent evolutionary experiment, testing for the consistency of the relationship’s direction rather than its numerical magnitude. This is crucial because the absolute values of speciation, extinction, and transition rates may not be directly comparable across clades with vastly different ages, sizes, and unique evolutionary histories [53]. Furthermore, this method is robust to the “label switching” issue inherent in hidden Markov models, where “rate class A” in one clade is not necessarily equivalent to “rate class A” in another. Because our test only considers the internal correlation, a positive relationship is recorded regardless of whether it is rate class A or B that has the faster rates. Under the null hypothesis of no consistent association, we would expect a random mix of positive and negative relationships across the clades, yielding a median of zero.

## Results and Discussion

### Biome shift rates are positively correlated with turnover rates

The colonization of new biomes is frequently hypothesized to be a major driver of diversification, with some studies showing that geographic movement into new areas can be more tightly linked to diversification than morphological innovation [53,54]. However, this relationship is known to be complex and context-dependent, as not all dispersal events trigger diversification bursts [55–57]. These contingent results suggest that the unique history of each clade plays a significant role in shaping its evolutionary trajectory, potentially obscuring more general patterns. A promising way to move beyond these clade-specific effects is to examine the dynamics within major clades, treating each as an independent evolutionary experiment. This approach allows for a more focused test of whether the intrinsic lability of a lineage (its propensity to transition between biomes) is generally associated with a faster tempo of diversification, rather than focusing on the impact of singular biome shifts.

Our analysis reveals a consistent, positive relationship between the rate of biome shifts and the rate of evolutionary turnover across angiosperm clades, but no corresponding association with net diversification. Using a non-parametric sign test, we found that the rate class with higher biome transition rates was significantly more likely to also have higher rates of turnover (speciation + extinction). The median of the slopes describing this relationship was positive (0.23), and the 95% confidence interval for the median [0.00, 4.41] excludes negative values, providing statistical support for a consistent positive association (p = 0.016). In contrast, we found no consistent directional relationship between biome shift rates and net diversification rates (speciation - extinction). For this comparison, the median slope was zero, with a wide 95% confidence interval [0.00, 6.48], indicating a lack of any consistent directional trend (p = 0.125).

These patterns are evident in the parameter estimates from the best-fit model for each of the 13 clades analyzed (Table 1; Table S3, S4). All of the clades showed support for hidden rate classes and the majority (7 of 13) showed a positive association between transition rates and turnover rates (Figure 2). This pattern was particularly strong in clades like Fagales, Onagraceae, and Arecaceae, where the rate class with more frequent biome shifts also exhibited substantially higher turnover. The case of Onagraceae is particularly interesting because it possessed a strong positive association between biome shifts and turnover, but a strongly negative association with net diversification (mTurn = 8.637, mNetDiv = −501.167). While the extreme magnitude of the net diversification slope should be interpreted with caution (the potential for high variance in parameter estimates is why a non-parametric test was used here), the opposing direction of the two relationships is interesting. The Onagraceae result suggests a scenario where higher lability is associated with higher turnover (more macroevolutionary events), but this increased turnover is associated with lower net diversification (less lineages surviving to the present). In other words, for this clade, the state characterized by frequent biome shifts accumulates species at a slower pace than the more stable state and the increase in speciation rates associated with lability is less than the corresponding increase in extinction.

**Table 1.** Parameter estimates from the best-fit hidden Markov models. For each of the 13 clades, the table shows the Akaike weight (AICwt) of the best-fit model and the estimated mean rates for biome transitions (meanTrans), turnover (meanTurn), and net diversification (meanNetDiv) for the two hidden rate classes (A and B). The slope columns (slopeTurn, slopeNetDiv) represent the relationship between the rate of biome shifts and diversification rates. A slope of zero indicates that the best-fit model constrained transition rates to be equal across the hidden states.

| clade | nTip | AICwt | meanTransA | meanTurnA | meanNetDivA | meanTransB | meanTurnB | meanNetDivB | slopeTurn | slopeNetDiv |
| --- | --- | --- | --- | --- | --- | --- | --- | --- | --- | --- |
| Fagales | 526 | 0.898 | 0.076 | 0.322 | 0.093 | 28.834 | 1.382 | 0.4 | 27.132 | 93.844 |
| Onagraceae | 267 | 0.768 | 39.369 | 4.802 | 0.026 | 0.007 | 0.245 | 0.105 | 8.637 | -501.167 |
| Arecaceae | 2202 | 0.992 | 1.76 | 0.367 | 0.367 | 0.169 | 0.104 | 0.104 | 6.032 | 6.032 |
| Arabideae | 320 | 0.571 | 0.163 | 0.752 | 0.051 | 9.783 | 5.789 | 0.393 | 1.909 | 28.156 |
| Carex | 1221 | 0.798 | 0.336 | 0.281 | 0.281 | 0.596 | 0.582 | 0.582 | 0.861 | 0.861 |
| Salvia | 484 | 0.816 | 0.073 | 0.132 | 0.105 | 0.167 | 0.515 | 0.409 | 0.246 | 0.31 |
| Eucalypts | 691 | 1 | 2.081 | 9.035 | 0.302 | 0.129 | 0.39 | 0.013 | 0.226 | 6.765 |
| Antirrhineae | 276 | 0.194 | 0.453 | 0.167 | 0.167 | 0.453 | 0.458 | 0.458 | 0 | 0 |
| Apioideae | 1033 | 0.938 | 0.934 | 1.187 | 0.378 | 0.934 | 0.55 | 0.163 | 0 | 0 |
| Mimosoids | 1758 | 1 | 0.367 | 0.161 | 0.133 | 0.367 | 0.416 | 0.35 | 0 | 0 |
| NeoAstragalus | 364 | 0.997 | 34.923 | 0.725 | 0.725 | 34.923 | 0.162 | 0.162 | 0 | 0 |
| Restionaceae | 332 | 0.402 | 0.081 | 0.027 | 0.027 | 0.081 | 0.122 | 0.122 | 0 | 0 |
| Solanaceae | 1000 | 1 | 0.457 | 0.432 | 0.207 | 0.457 | 1.105 | 0.529 | 0 | 0 |

**Figure 2.**
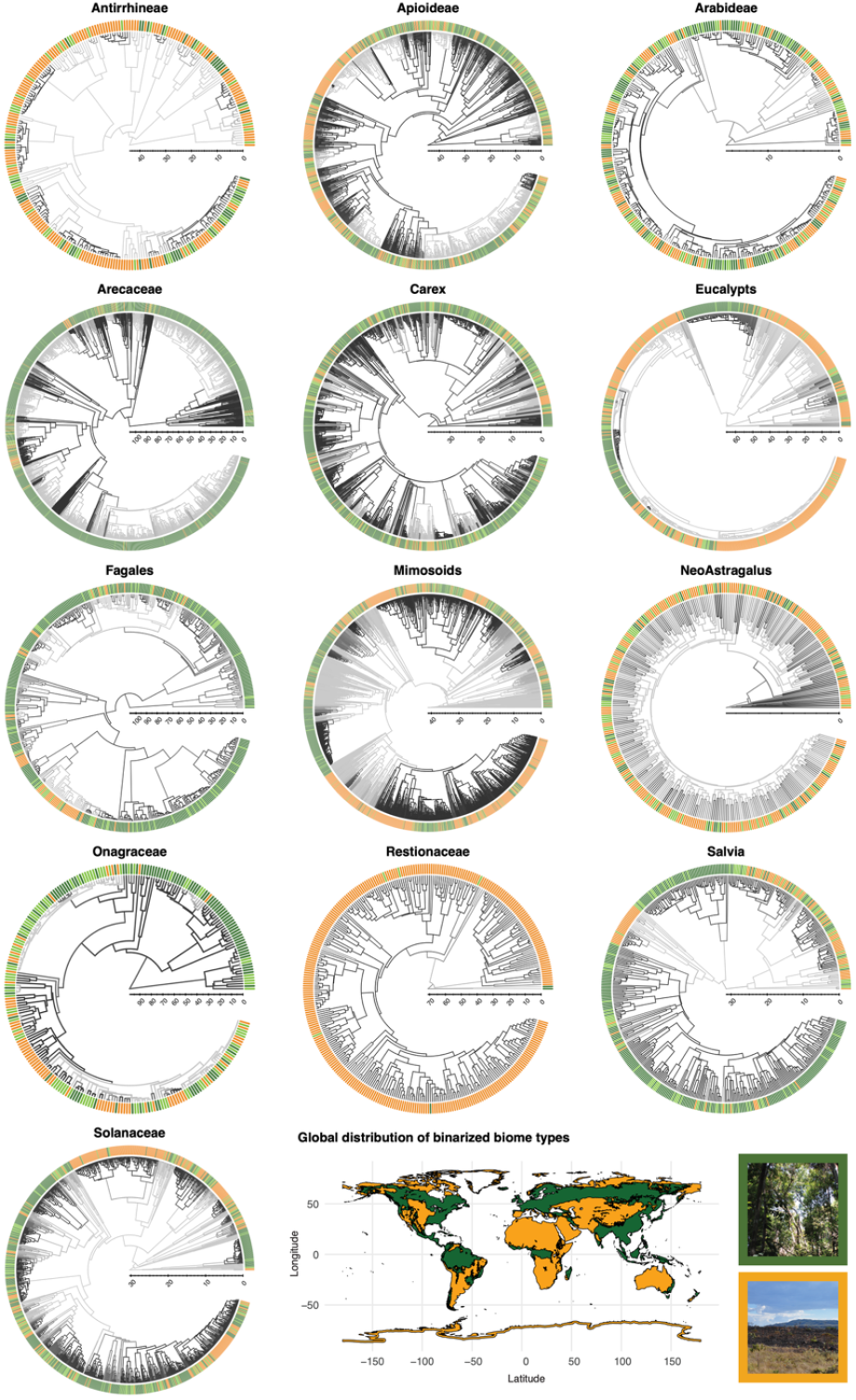
Phylogenetic distribution of biome affiliation and rate-class heterogeneity across 13 angiosperm clades. Each circular phylogeny represents a single focal clade, with tips color-coded according to biome association: closed-canopy (dark green), open-canopy (orange), or widespread (light green). Phylogenetic trees are scaled by evolutionary time, and internal branching patterns illustrate inferred diversification dynamics from the best-fitting hidden-state speciation and extinction (HiSSE) models. The bottom panel shows the global distribution of binarized biome types (closed-versus open-canopy) used to score species occurrences based on GBIF records and WWF biome classifications. Photographs depict representative closed-(top) and open-canopy (bottom) habitats.

Finally, for six of the thirteen clades (e.g., Mimosoids, Solanaceae, NeoAstragalus), the best-fit models constrained biome transition rates to be equal across the hidden rate classes (Figure 2; Table 1). In these instances, our analysis by definition could not detect a relationship between lability and diversification, resulting in a slope of zero. However, the lack of any negative associations across the remaining clades is informative. In every case where a relationship could be assessed, higher rates of biome shifts were associated with higher—never lower—rates of turnover. This suggests that while the strength of this coupling may vary, or may not be detectable in all lineages, we found no evidence that increased lability ever acts to slow the pace of evolution. Taken together, our results support the hypothesis that lability in biome preference has red-queen type dynamics [58], elevating the pace of both speciation and extinction, but it is not a consistent predictor of long-term species accumulation.

### An alternative perspective on diversification analyses

State dependent diversification analyses (SSE models) began as a way to correct biases in transition rates due to the possibility that observed states may be correlated with speciation and extinction rates [18]. It was later discovered that this class of models was biased towards finding associations between a focal character and diversification rates [59]. This was because SSE models included heterogeneity in diversification parameters that would outperform alternative models which did not, regardless of whether there was a true association between diversification rates and the focal character [60]. Hidden Markov models, which were introduced as a more biologically plausible model of evolution by allowing for rate heterogeneity independent of the focal character (e.g. HiSSE, [9]) were able to be utilized as better null hypotheses and thus overcame challenges of previous SSE models (see also [61]). This body of work set the stage for the typical diversification analysis in which biologists test for an association between the focal character and diversification parameters with the inclusion of a hidden Markov model as a null hypothesis. More recently, it has also been suggested that these models are also relatively robust to noted identifiability issues of birth-death models [60].

Although treating hidden Markov SSE models as null hypotheses is common practice, there is a particularly powerful way in which these models have been underutilized. Instead of treating hidden states merely as a nuisance variable to be accounted for, the different rate classes inferred by these models can themselves be treated as the “trait” of interest. This approach allows for a direct test of the hypothesis that the lability of diversification rates—the propensity for a lineage to shift between different rates of speciation and extinction—is itself an important factor in explaining broad-scale patterns of diversity. Hidden Markov models, therefore, are not just null models against which to test trait-dependent hypotheses; they also offer a framework for exploring the role of evolutionary lability in shaping diversification dynamics.

HMMs are explicitly parameterized by the dynamics of the observed characters, as this is where the information for estimating rate classes is derived from [20]. This means that evidence for hidden Markov models can come from both differences in diversification dynamics and differences in observed character transition rates, and the potential relationships between these processes has been generally under-explored. This is not to say that biologists have ignored the potential for trait lability to drive patterns of diversification. Several studies have attempted to correlate these processes using independent modeling results (e.g. [8,62]), but not in a framework where the diversification and trait evolution parameters were estimated jointly. Furthermore, unlike many studies which utilize SSE models, we are not primarily interested in whether a particular character influences diversification. The hidden Markov SSE models allow us to frame our hypothesis in such a way that the rate classes themselves are of primary interest. This opens up potential examinations of other rate class dynamics as the underlying character evolution model can be structured to match different biological hypotheses. In our case, we are testing whether the rates of biome shifts are positively correlated with rates of net-diversification and turnover in a joint framework, but the tempo of the models is just one aspect of discrete character evolution.

Finally, this approach offers the potential to better account for the clade-specific differences. By modeling rate heterogeneity within each major clade, it is possible to test for a consistent relationship between state transitions (e.g., biome shifts) and diversification rates without the unique, contingent histories of those clades acting as confounding factors. In this way, each clade can be treated as its own evolutionary experiment, providing independent tests of the hypothesis that higher rates of trait transition are correlated with faster rates of diversification. The framework outlined here can be implemented using existing tools as it does not require any new methodological developments. Instead, it relies on evolutionary biologists to refocus their hypotheses in an examination of how different processes influence diversification rather than particular character traits.

## Conclusion

The significant relationship between rates of biome shifts and turnover highlights the role that trait lability can play in shaping the diversification dynamics across many clades. Though lability is not a given for all clades, when it is present it may be a signature of important biological shifts which are reflected in both diversification and trait evolution. This fact highlights one of the main methodological differences of our study. Rather than using SSE models and hidden Markov SSE models to test whether a particular phenotype influences diversification, we examine the possibility that diversification dynamics are influenced by the processes underlying phenotypic evolution, the tempo of biome change in this case. This possibility was recognized by G. Ledyard Stebbins 50 years ago and proposed by several others [e.g., 8,11] to be one of the primary reasons for the high species diversity in angiosperms. However, this conclusion does not entirely bear out in our results because net-diversification, the parameter related to species accumulation, is not significantly tied to the tempo of biome shifts. Instead, biome lability may be a double-edged sword increasing the propensity of lineage speciation, but equally increasing extinction risk.

## Supplemental Material

**Table S1.** This table outlines the structure of the 36 unique models tested for each angiosperm clade. Each row represents a distinct model defined by the combination of parameter settings for four primary components: extinction fraction (ef), turnover (turn), the transition rate matrix (q), and the rate class structure (rc). The number associated with each component (e.g., ef1, ef2) indicates a specific parameterization that dictates how rates are allowed to vary. For instance, ef1 corresponds to a model with no variation in the extinction fraction across states. The 36 models represent the full set of combinations of these different configurations for extinction, speciation, and biome transition dynamics.

| ef | turn | q | rc |  |  |  |
| --- | --- | --- | --- | --- | --- | --- |
| 1 | 1 | 1 | 1R | **model class** | **parameters to vary (ef)** | **parameters to vary (turn)** |
| 2 | 1 | 1 | 1R | CID | ef1 = 0,1,1,1,0,1,1,1 | turn1 = 0,1,1,1,0,1,1,1 |
| 1 | 2 | 1 | 1R | CD | ef2 = 0,1,2,3,0,1,2,3 | turn2 = 0,1,2,3,0,1,2,3 |
| 2 | 2 | 1 | 1R | CID | ef3 = 0,1,1,1,0,2,2,2 | turn3 = 0,1,1,1,0,2,2,2 |
| 1 | 1 | 1 | 2R | CD | ef4 = 0,1,2,3,0,4,5,6 | turn4 = 0,1,2,3,0,4,5,6 |
| 2 | 1 | 1 | 2R |  |  |  |
| 3 | 1 | 1 | 2R |  | **parameters to vary (q)** | **parameters to vary (rc)** |
| 4 | 1 | 1 | 2R |  | q1 = no difference between rate classes | 1R = one rate class |
| 1 | 2 | 1 | 2R |  | q2 = difference between rate classes | 2R = two rate class |
| 2 | 2 | 1 | 2R |  |  |  |
| 3 | 2 | 1 | 2R | **example** | **description** |  |
| 4 | 2 | 1 | 2R | ef1,turn1,q1,1R | dull null. no state dependent differences. |  |
| 1 | 3 | 1 | 2R | ef2,turn2,q1,1R | bisse. classic state dependent diversification. |  |
| 2 | 3 | 1 | 2R | ef3,turn3,q1,1R | CID2. null model with hidden states |  |
| 3 | 3 | 1 | 2R | ef3,turn3,q2,1R | null model w.r.t. diversification, but allowing different trans rates |  |
| 4 | 3 | 1 | 2R |  |  |  |
| 1 | 4 | 1 | 2R |  |  |  |
| 2 | 4 | 1 | 2R |  |  |  |
| 3 | 4 | 1 | 2R |  |  |  |
| 4 | 4 | 1 | 2R |  |  |  |
| 1 | 1 | 2 | 2R |  |  |  |
| 2 | 1 | 2 | 2R |  |  |  |
| 3 | 1 | 2 | 2R |  |  |  |
| 4 | 1 | 2 | 2R |  |  |  |
| 1 | 2 | 2 | 2R |  |  |  |
| 2 | 2 | 2 | 2R |  |  |  |
| 3 | 2 | 2 | 2R |  |  |  |
| 4 | 2 | 2 | 2R |  |  |  |
| 1 | 3 | 2 | 2R |  |  |  |
| 2 | 3 | 2 | 2R |  |  |  |
| 3 | 3 | 2 | 2R |  |  |  |
| 4 | 3 | 2 | 2R |  |  |  |
| 1 | 4 | 2 | 2R |  |  |  |
| 2 | 4 | 2 | 2R |  |  |  |
| 3 | 4 | 2 | 2R |  |  |  |
| 4 | 4 | 2 | 2R |  |  |  |

**Table S2.**
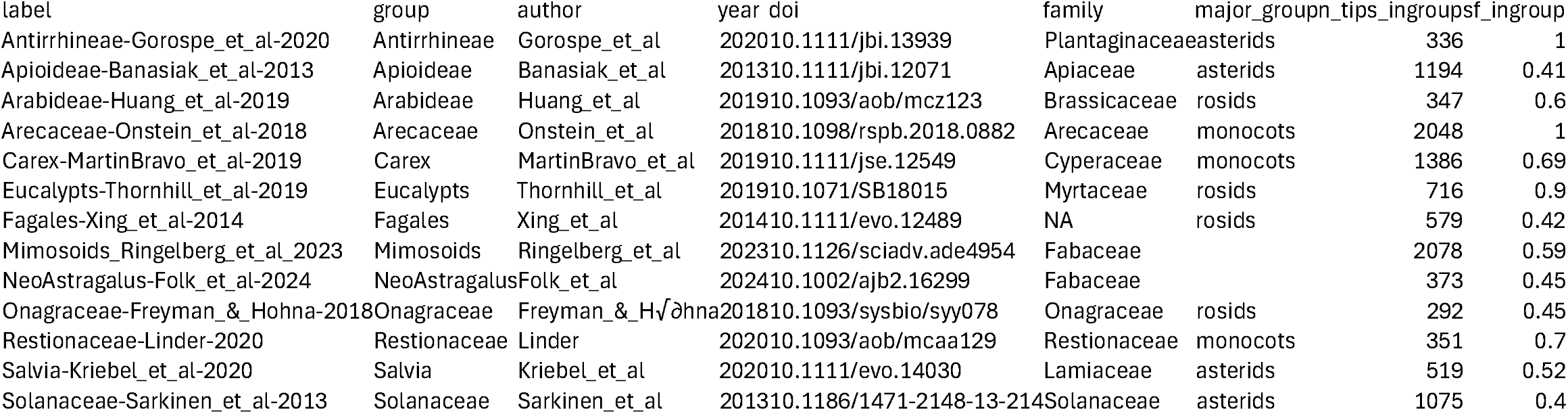
Sampling fractions for each clade.

**Table S3.**
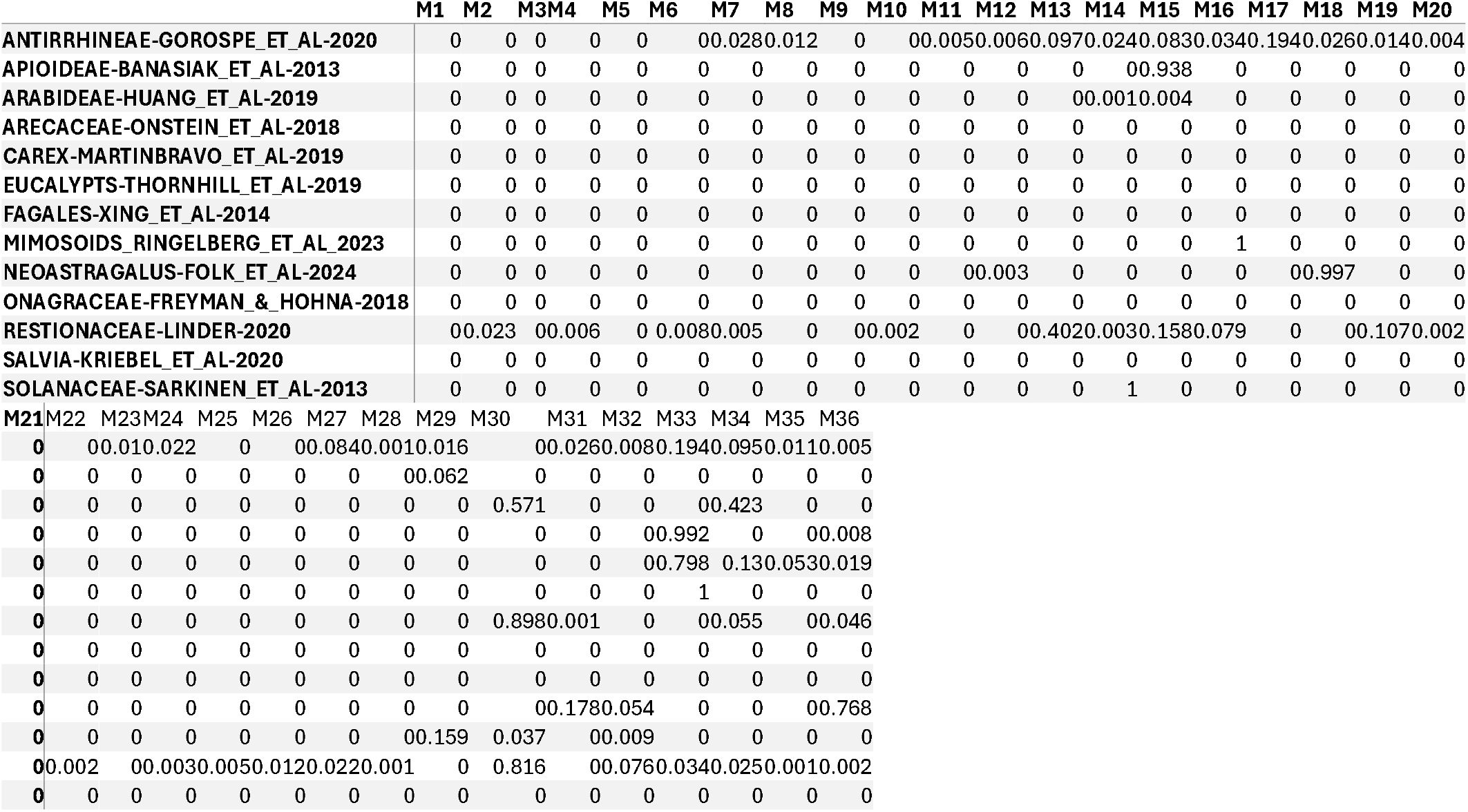
AIC weights for all 36 model fits for each clade. See Table S1 for the meaning of each model (M1 to M36).

**Table S4.**
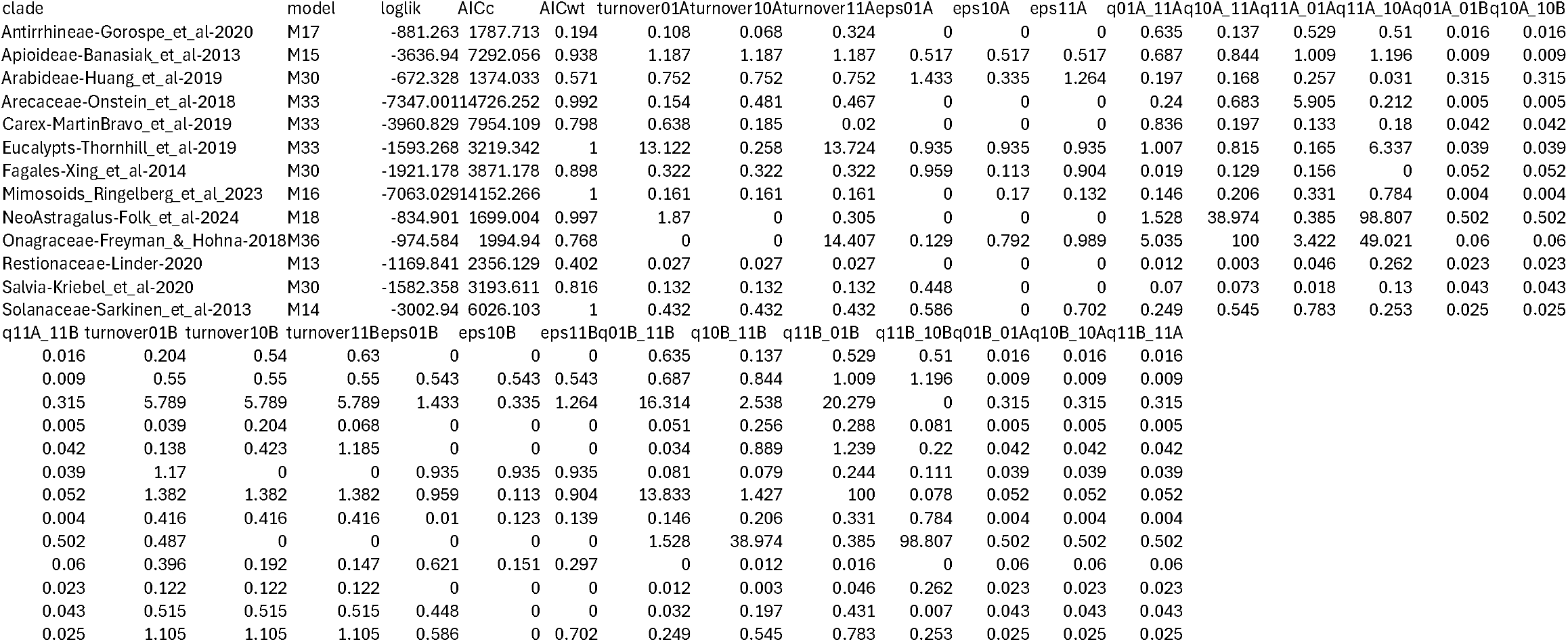
Parameter estimates for each of the best fitting models.

